# ReFeaFi: Genome-wide prediction of regulatory elements driving transcription initiation

**DOI:** 10.1101/2021.03.31.437992

**Authors:** Ramzan Umarov, Yu Li, Takahiro Arakawa, Satoshi Takizawa, Xin Gao, Erik Arner

**Affiliations:** Graduate School of Integrated Sciences for Life, Hiroshima University, Higashi-Hiroshima, 739-8528 Japan; Department of Computer Science and Engineering (CSE), The Chinese University of Hong Kong (CUHK), Hong Kong, People’s Republic of China; Laboratory for Applied Regulatory Genomics Network Analysis, RIKEN Center for Integrative Medical Sciences, Yokohama, Kanagawa, 230-0045 Japan; King Abdullah University of Science and Technology, Computational Bioscience Research Center, Computer, Electrical and Mathematical Sciences and Engineering Division, Thuwal 23955-6900, Saudi Arabia

## Abstract

Regulatory elements control gene expression through transcription initiation (promoters) and by enhancing transcription at distant regions (enhancers). Accurate identification of regulatory elements is fundamental for annotating genomes and understanding gene expression patterns. While there are many attempts to develop computational promoter and enhancer identification methods, reliable tools to analyze long genomic sequences are still lacking. Prediction methods often perform poorly on the genome-wide scale because the number of negatives is much higher than that in the training sets. To address this issue, we propose a dynamic negative set updating scheme with a two-model approach, using one model for scanning the genome and the other one for testing candidate positions. The developed method achieves good genome-level performance and maintains robust performance when applied to other species, without re-training. Moreover, the unannotated predicted regulatory regions made on the human genome are enriched for disease-associated variants, suggesting them to be potentially true regulatory elements rather than false positives. We validated high scoring “false positive” predictions using reporter assay and all tested candidates were successfully validated, demonstrating the ability of our method to discover novel human regulatory regions.

## Introduction

The study of gene regulation is primarily concerned with two classes of regulatory elements: promoters, which define the Transcription Start Site (TSS), and enhancers, that amplify the transcription^1^. The TSS is the first nucleotide that is copied at the 5’ end of the corresponding mRNA. A core promoter is a minimal promoter region that typically spans several hundred bp up- and downstream of a TSS and is capable of initiating basal transcription^2^. Core promoters have complex and gene-specific architectures consisting of unique compositions of binding sites for Transcription Factors (TFs) involved in specific regulation of transcription. Transcription is further stimulated by enhancer elements, which can be located at a long distance from the target core promoter. These distal locations are able to affect the transcription due to a favorable folding of the genome in the three-dimensional space^3^. Enhancer sequences are also capable of bidirectional transcription of RNAs (eRNAs) at a large scale. This means that gene promoters and enhancers share a similar promoter architecture, each bound by RNA pol II when active^4^. Recent studies have shown that promoters and enhancers share several properties and functions related to their chromatin and sequence architectures^5^. The distinction between these regulatory elements is further reduced by the fact that promoters can enhance transcription at a distant site^6^ while enhancers can drive local transcription initiation^7^. This provides a motivation to consider these elements together when trying to understand transcription regulation and build models for their identification.

Thanks to the development of advanced experimental techniques, significant progress has been made in identifying gene regulatory sequences^8–10^ However, a detailed experimental exploration of transcripts is still an expensive and difficult procedure. Therefore, in addition to experimental efforts, accurate computational identification of putative regulatory regions, for both individual genes and entire genomes, remains an important challenge of genomics studies. Computational prediction is important for guiding experimental biologists, providing putative regions which can be validated using reporter gene assays.

Accurate computational identification of regulatory elements remains a difficult task due to the high diversity of DNA sequence features and tissue specificity of the transcriptional regulation. There are numerous computational tools developed in an attempt to predict promoters and enhancers^11–14^. However, many of them focus on discrimination between fixed sets of promoter/enhancer sequences and random genomic sequences. The reported performance deduced from such small and balanced test sets does not hold when evaluated at the level of the whole genome, which is a much more difficult task due to the huge number of tested locations^15^.

It has been shown that accurate regulatory element prediction can be achieved on the genome wide scale by using epigenetic data, such as DNA methylation and histone modification profiling^16^. For example, H3K4me1, H3K4me3, and H3K27ac marks are associated with promoter and enhancer activities^17^. In recent years, a number of methods have been developed that utilize local epigenetic marks for regulatory element prediction, based on machine learning algorithms such as random forests^16,18^, support vector machines^19,20^, and deep learning^21,22^. However, there is still a need for methods that can provide accurate predictions based on DNA sequences, in particular for annotating the genomes of species where epigenetic data are not widely available.

In this study, we developed ReFeaFi (Regulatory Feature Finder), a general genome-wide promoter and enhancer predictor, using the DNA sequence alone. Using Cap Analysis Gene Expression (CAGE) data^23^ as the ground truth for promoters and enhancers, we used a dynamic training set updating scheme to train the deep learning model, which allows us to have high recall while keeping the number of false positives low, improving the discrimination and generalization power of the model. ReFeaFi achieved comparable performance when the model trained on the human genome was applied to other vertebrate species, showing the generality of our model. We found that unannotated regulatory regions predicted by our method are enriched for genetic variants associated with disease^24,25^, suggesting that they might be real regulatory elements. High scoring unannotated predictions were validated using reporter assays and all the candidates showed strong luciferase signal. By analyzing synthetic promoters, we found that the predicted score strongly correlates with measured expression strength. We used the trained deep learning model to study the architecture of regulatory elements and to find out how conserved elements affect transcription strength. Finally, we have developed a novel model analysis technique that reveals related positions around the TSS.

## Results

### Genome-wide identification of regulatory elements

The overview of our method is shown in **Figure 1**. The method is two-tiered and consists of a scan model and a prediction model, which are trained iteratively to reduce false positives (FPs). Briefly, the scan model picks candidate regions for the prediction model, which then makes a decision if these regions contain one or more TSS. If an unannotated region receives a score above the threshold, it is added to the negative set. The whole process is repeated several times to generate a difficult negative set which forces the model to learn more complex features for the identification of regulatory regions. We trained our model on the human genome, with chromosome 1 used as the test set and the rest of the genome for training and validation. The positive set of regulatory region annotations used was constructed by merging the human robust promoter set and human permissive enhancer set downloaded from the FANTOM5 website^26^. We initially compared our method against several previously published promoter predictors (Basenji^27^, PromPredict^28^, and EP3^11^), which are capable of genome-wide TSS predictions. Using human chromosome 1 for evaluation, our developed model achieved good performance, significantly outperforming other methods (**Table 1, Figure 2a**). In particular, ReFeaFi generated substantially less FPs than a recently proposed deep learning-based method for regulatory elements prediction (Basenji), for all recall values, see **Figure 2a**.

**Figure 1.**
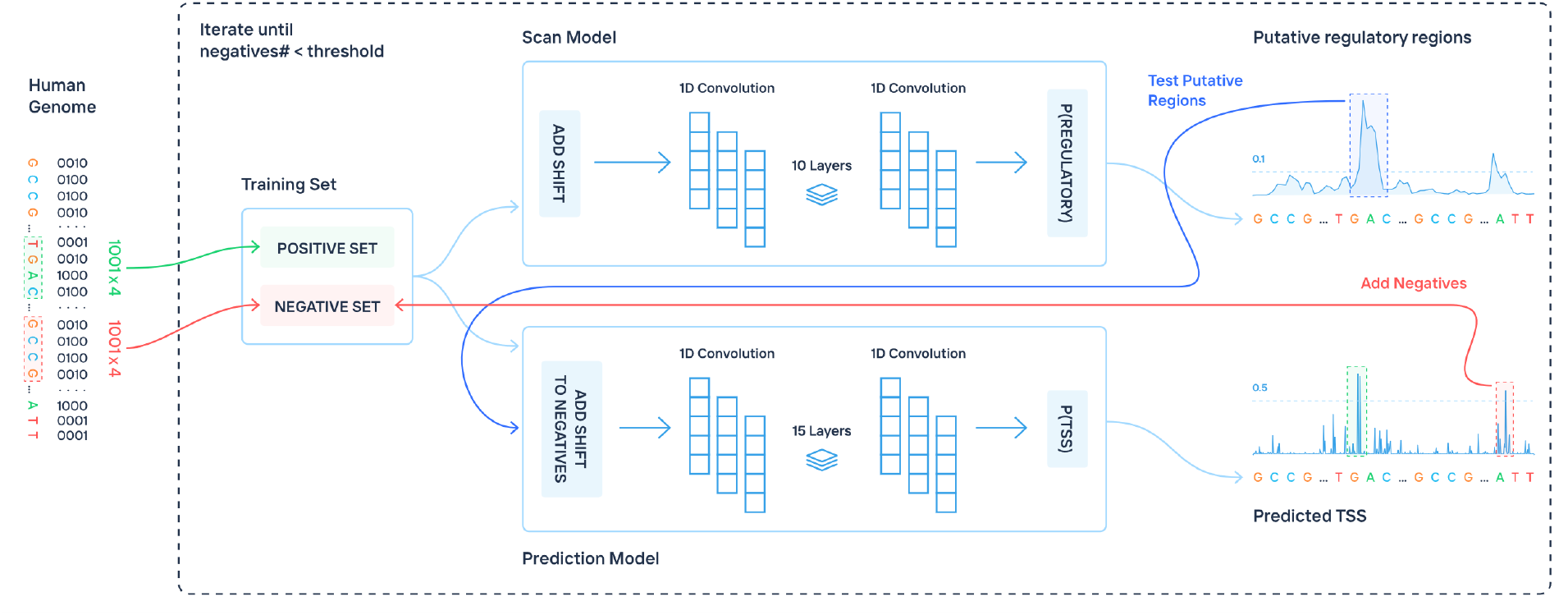
Workflow of the proposed method for genome-wide regulatory elements prediction. The scan model uses a sliding window approach to pick putative regulatory regions. The prediction model finds TSS positions inside these regions by testing each position. The false positive predictions made by the second model are added to the negative set for the next round of training. The whole process is repeated iteratively to generate a difficult negative set which forces the model to learn how to distinguish the difficult negatives from the real regulatory sequences.

**Table 1.** Comparison of the performance of different TSS identification methods. The decision thresholds for these methods were adjusted to achieve the same recall of 0.50.

| Method | Recall | Precision | F1 score | FP per correct | FP per 1 Mb |
| --- | --- | --- | --- | --- | --- |
| ReFeaFi | 0.50 | 0.52 | 0.51 | 0.93 | 46.44 |
| EP3 | 0.50 | 0.20 | 0.29 | 4.01 | 198.26 |
| Basenji | 0.50 | 0.07 | 0.12 | 13.48 | 664.92 |
| PromPredict | 0.50 | 0.07 | 0.12 | 13.44 | 666.26 |

**Figure 2.**
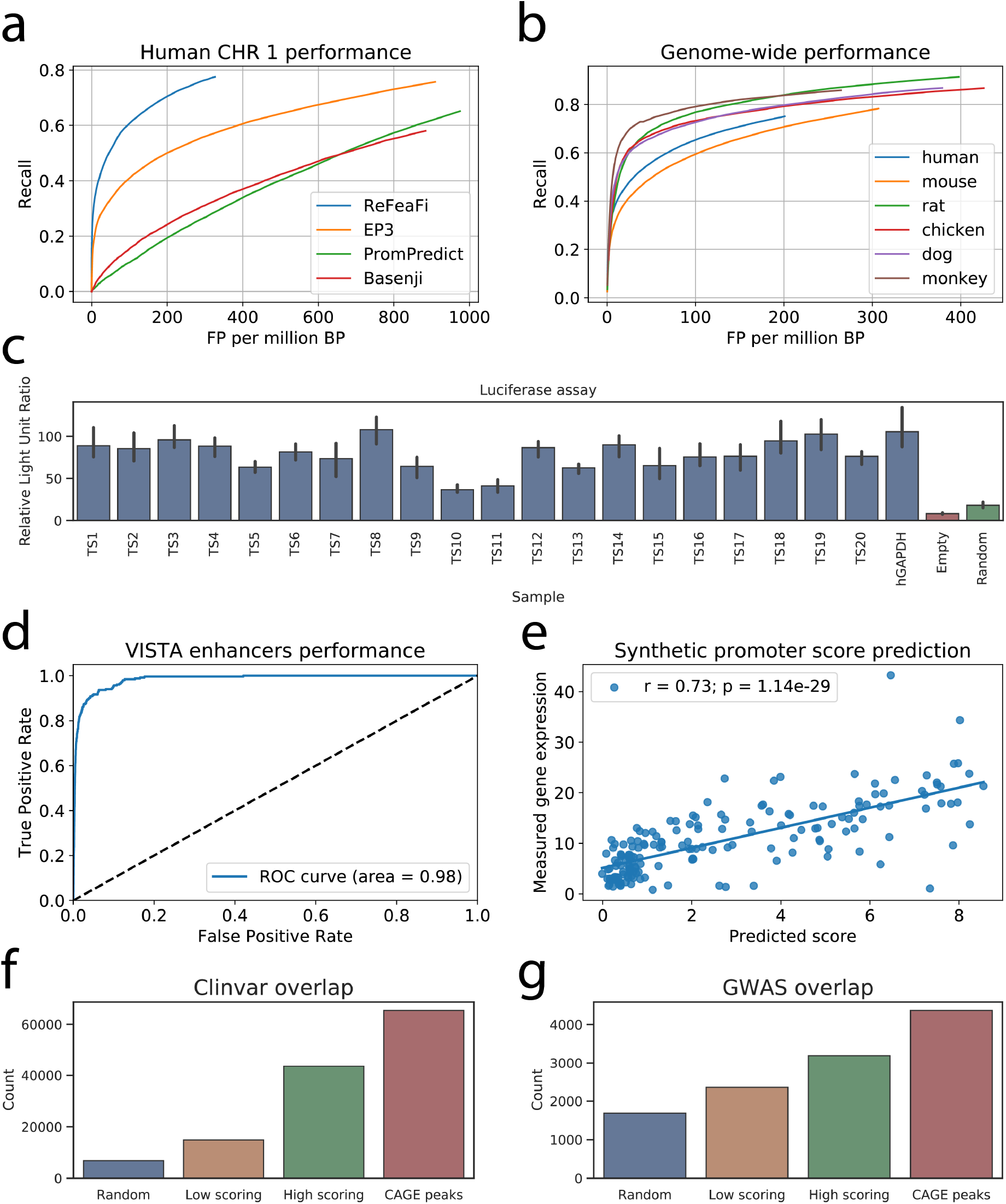
The performance evaluation of the proposed method. **a.** Performance of TSS predictors on human chromosome 1. Our method, ReFeaFi, significantly outperforms other predictors. **b.** Performance of the proposed method applied on different vertebrate species, without re-training the model. Due to the small number of known CAGE peaks, there is a significant difference between human/mouse and other organisms. **c.** Results of the reporter assay on the predicted unannotated regulatory regions. Each chosen sample is at least 5kb away from any known CAGE peaks or GENCODE annotations. Empty vector and random sequences were used as negative controls, while GAPDH promoter is used as positive control. All the candidates passed the validation, with several candidates showing intensity similar to the positive control. **d.** ReFeaFi achieves outstanding performance on discriminating VISTA enhancers and 100 times as many random genomic regions. **e.** Correlation between score predicted by our method and measured mean expression of the synthetic promoters. **f.** The number of Clinvar genetic variants located in the vicinity of low and high scoring predicted regions. **g.** GWAS genetic variants overlap with the predicted regions.

We subsequently applied the model trained on human CAGE data to genomes of other species, without re-training (**Figure 2b)**. The obtained result for the mouse genome was similar to the human performance, achieving comparable recall and FP rate. For other species, the recall was high, but the FP rate was significantly higher than for human and mouse. This is expected and due to the fact that the transcriptomes of these species have been profiled at considerably lower depth, and there are thus much fewer CAGE peaks known for these species (**Supplementary Table 1,** most of them are related to housekeeping genes which are easier to detect due to higher GC content^29^). This gives rise to a high recall, where additional FPs are most likely related to tissue-specific promoters, which have not been detected in the currently available CAGE datasets for these organisms.

### Validation of unannotated predicted regions by reporter assay

We next set out to investigate the extent to which predicted regulatory regions marked as false positives in the initial evaluation might represent regions with true regulatory potential, but not yet discovered by the current experimental annotations. To this end, we performed reporter assays of 17 high scoring regions and three regions with a low score but still predicted as regulatory by our model. The tested sequences included regions with and without classic promoter elements (INR and TATA box, **Supplementary Table 2**) and were chosen so that they were at least 5kb away from any known CAGE peaks or GENCODE annotations. All the chosen candidates showed significantly higher luciferase signals than negative controls (empty vector and random sequence, **Figure 2c**), with several sequences showing intensity similar to the GAPDH promoter used as the positive control.

### Highly accurate prediction of VISTA enhancers

The VISTA Enhancer Browser contains experimentally validated human and mouse enhancers with their activity measured in transgenic mice^30^. VISTA enhancers were previously used by Yang *et al*. to validate their enhancer predictor BiRen^13^, where like our method, the model is trained using DNA sequences alone, unlike methods that use epigenetic information to predict enhancers^19,20^. In this validation approach, a test set is constructed by using VISTA enhancers as a positive set, with a negative set that contains ten times as many non-enhancer sequences. When evaluated, BiRen achieved AUC of 0.945, improving over DEEP^14^ and SVM^31^ enhancer predictors which achieved AUC of 0.883 and 0.621, respectively. We adopted the same experimental setup, constructing a positive set with all the VISTA human and mouse enhancers from chromosome 1 and 10 times more random genomic regions for the negative set. Our model achieved almost perfect accuracy, with AUC > 0.99. The same result was obtained when repeating the experiment with 100 times more negative sequences (AUC 0.98, **Figure 2d**).

### Predicted score correlates with measured expression

Weingarten-Gabbay *et al*. devised a high-throughput assay to quantify the activity of fully designed sequences that were integrated and expressed from a fixed location within the human genome^32^, using the method to investigate binding regions of core promoters. We applied our model to the designed sequences and computed the correlation between the measured expression and our predicted scores (**Figure 2e**). Correlation between the score and measured expression was 0.73, further showing the generality of our model, since it had not seen any of these synthetic sequences during training and suggesting that our model may be useful for designing new promoters with desired strength by screening potential candidate sequences.

### Non-annotated predicted regions are enriched for disease-associated variants

Nucleotide variation in enhancers and promoter regions has been shown to be associated with human diseases. Indeed, most of the disease-associated genome-wide association studies (GWAS) hits fall in the non-coding regions of the human genome^33^. GWAS helps to understand disease mechanisms and provides the starting point for the development of medical diagnosis, prognosis, and treatments. We tested if our genomewide predictions are enriched for genetic variants from GWAS^33^ and Clinvar^25^ databases. First, we computed the overlap between the known CAGE peaks and GWAS and Clinvar sets with a margin of 200 bp relative to the TSSs. Next, we compared the overlap with the variant sets for the same number of low scoring and high scoring predicted regions, excluding predictions overlapping known TSS or enhancers. The Clinvar set overlapped the high scoring regions three times more compared to the low scoring regions (**Figure 2f**), suggesting that they represent novel regulatory elements. The GWAS disease-associated SNPs were overrepresented in the high scoring regions as well, overlapping them two times more than expected by chance, **Figure 2g**. Together with the experimental results, this shows that the unannotated predicted regulatory regions we detected might have regulatory potential to a significant degree, and that they may be useful for prioritization of further studies of disease-associated variants.

### Model analysis reveals known and novel regulatory features

Given the limitations of the biochemical assays used to identify and characterize core promoter elements, it has been difficult to assign a clear function to each of these elements^2^. By feeding in modified regulatory sequences to our deep learning model, we can study how each core promoter element fine-tunes the gene expression. To measure the effects of each known motif on the promoter score, we searched for them using previously constructed PWMs for core promoter elements^34^ and replaced them with random nucleotides. New promoter scores were computed for the modified sequences and the change in the predicted score was recorded for each motif. The most important motifs were TATA-box and INR with 35% and 9% effect on the score, respectively (**Table 2**). Mutation of DPE and CCAT motifs had no strong effect on the predicted score, suggesting that they do not play a significant role in regulating human promoter activity. The same observations were made in a recent human promoter study^32^. In fact, the order in terms of gene expression impact is the same for the motifs we analyzed.

**Table 2.**
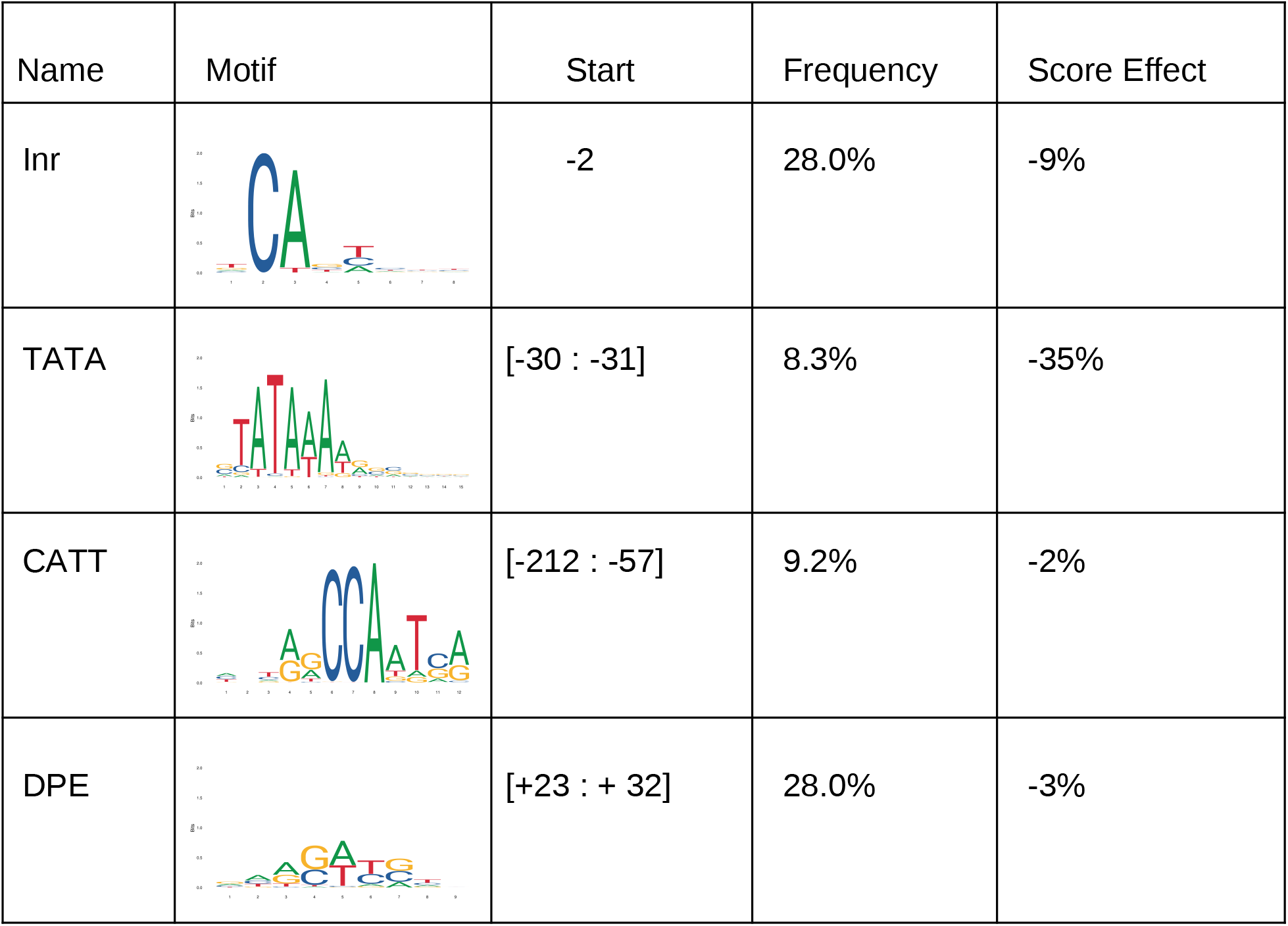
Effect of substituting promoter conserved motifs with random nucleotides on the score predicted by the deep learning model.

| Name | Motif | Start | Frequency | Score Effect |
| --- | --- | --- | --- | --- |
| Inr |  | -2 | 28.0% | -9% |
| TATA |  | [-30 : -31] | 8.3% | -35% |
| CATT |  | [-212 : -57] | 9.2% | -2% |
| DPE |  | [+23 : + 32] | 28.0% | -3% |

We next used the model to identify important locations in the input sequences that contribute most to the predicted score. Using a sliding window moving from the beginning of a regulatory sequence, we built a performance profile that reflects an effect of a random sequence, inserted in each sequence position in place of an original sequence, on the predicted score. The results for promoter and enhancer sequences (shown in **Figure 3a** and **Figure 3b** respectively) revealed that for promoters, the TSS and TATA regions are extremely important, while for enhancers, TSS regions contribute less to the total score compared to other regions.

**Figure 3.**
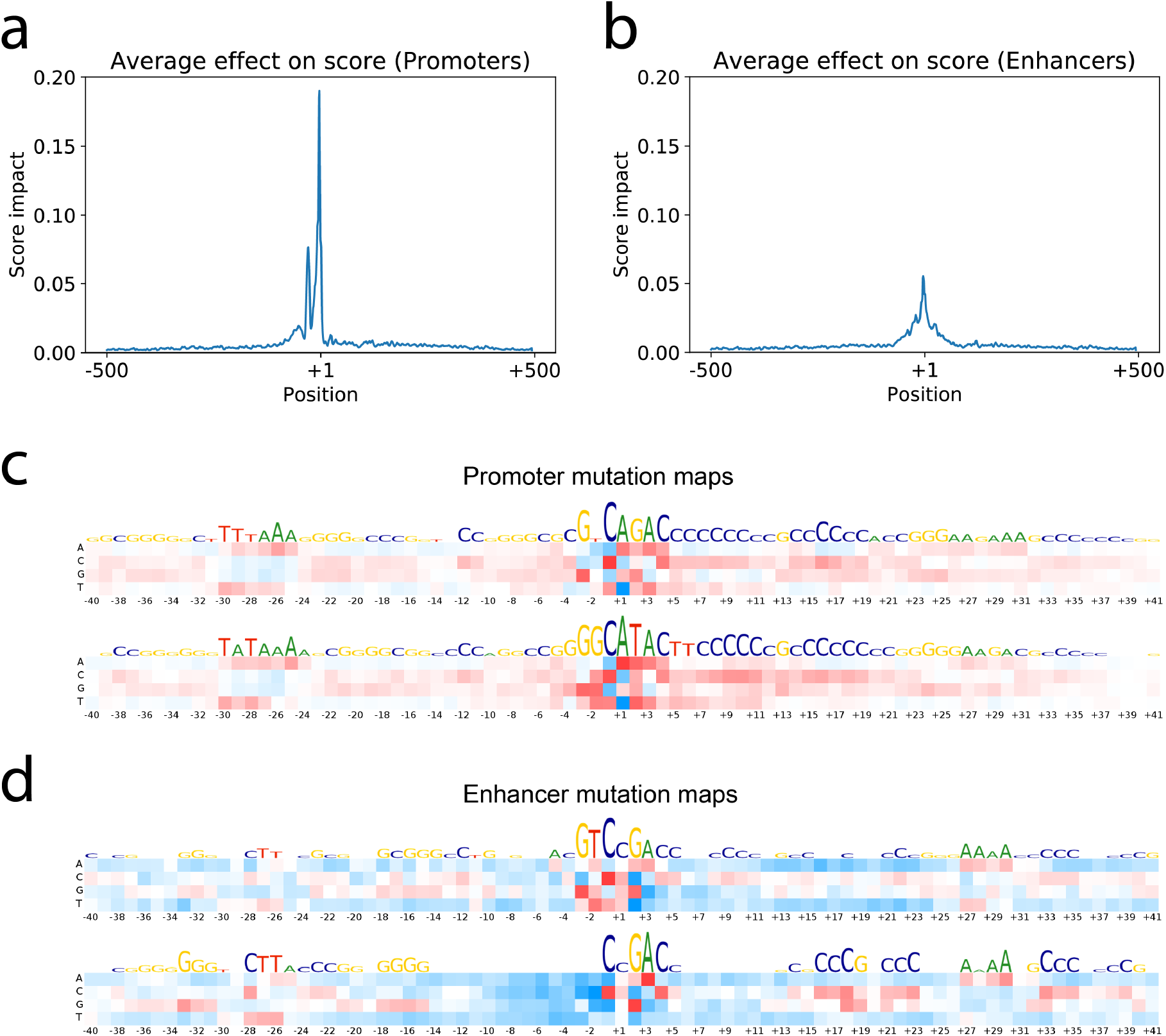
Analysis of the trained model. **a.** Effect of 7 bp sequence window substitution by a random sequence on the predicted score of the promoters. **b.** The same substitution procedure applied to the enhancer sequences. **c.** Mutation maps for the promoter sequences. K-means with k = 10 was applied to the mutation matrices before averaging and the two biggest clusters are shown. **d.** Mutation maps for the two biggest clusters of enhancer mutation matrices.

Despite the main effect on score coming from the core promoter region, in some cases replacing part of the sequences outside core promoter had a significant effect on the score, reducing it up to 50%. Motifs within core promoters have been extensively studied, but not much is known about distant motifs important for promoter activity. We tested if transcription factor binding motifs from the JASPAR database^35^ have a significant effect on the predicted score, and observed that even though there was often a perfect match of a known binding motif inside the promoter region, replacing it with random nucleotides did not change the output of the model in most cases. However, for some of the motifs, replacement with random nucleotides reduced the score significantly, also often when they were located far away from the TSS. Interestingly, top motifs for both promoter and enhancer sequences were the same, suggesting that they are regulated by a similar set of TFs. The TF binding motifs that were predicted to be the most important by our deep learning model are shown in **Table 3**. We only measured their effect on the predicted score when they were outside the core promoter region and still, they reduced the score by 22% on average in promoters and 19% in enhancers. We have also identified motifs that caused a significant decrease in the predicted score, however, did not have a match in the JASPAR database. The motifs were extracted and clustered based on their position relative to the TSS. The most influential motifs from the upstream and the downstream regions are shown in **Supplementary Table 3**.

**Table 3.**
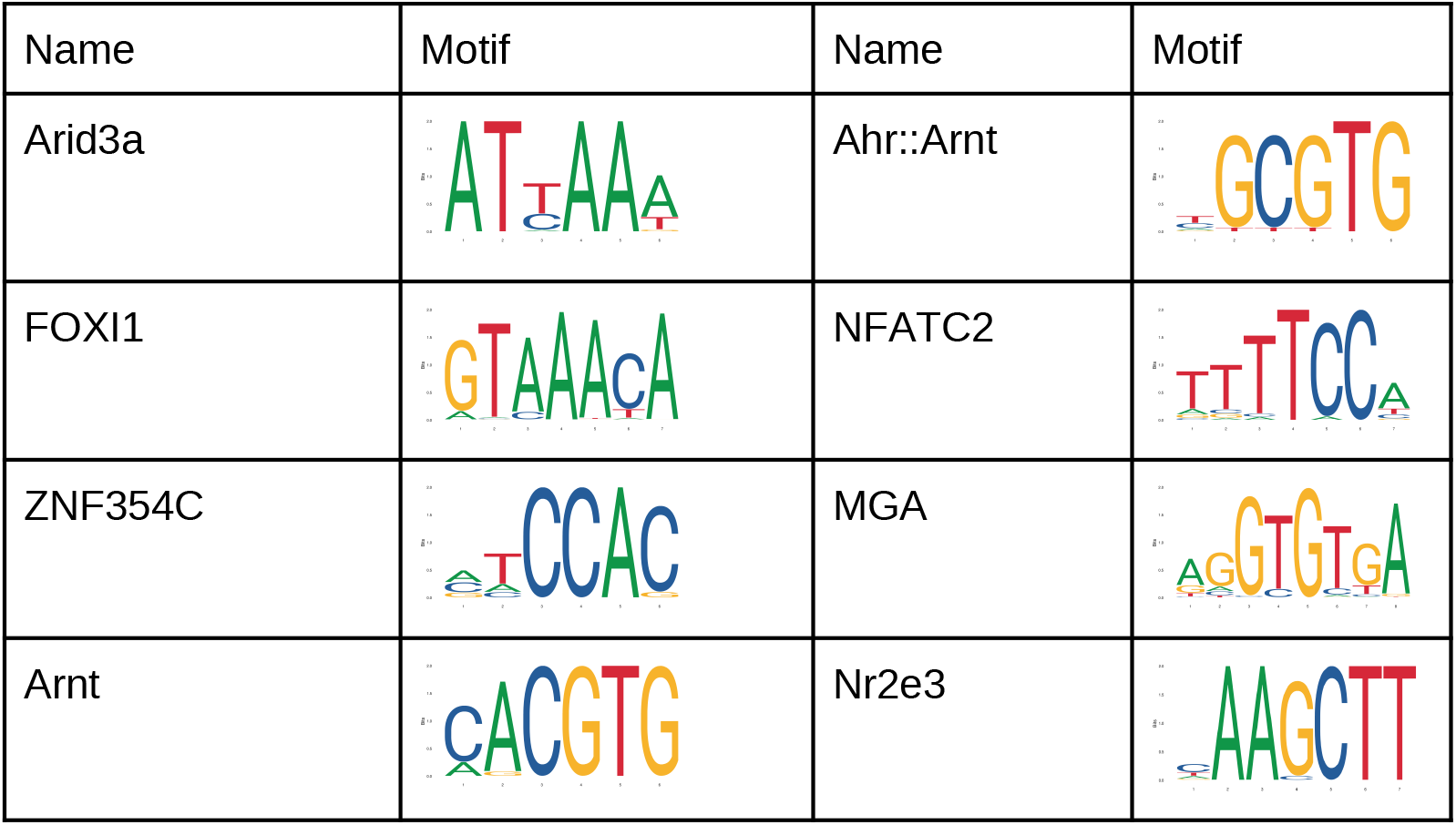
Most influential TF-binding motifs outside the core promoter region identified by our model for promoter and enhancer sequences.

| Name | Motif | Name | Motif |
| --- | --- | --- | --- |
| Arid3a |  | Ahr::Arnt |  |
| FOXI1 |  | NFATC2 |  |
| ZNF354C |  | MGA |  |
| Arnt |  | Nr2e3 |  |

To assess the contributions of different nucleotides in different positions of the regulatory sequences, we employed a modification of a feature mutation map for our test set. **Figure 3c** shows the mutation maps for the core promoter in the promoter sequences, while the mutation maps for the enhancer sequences is shown in **Figure 3d**. To build these maps, we studied how nucleotide substitutions will change the output score computed by our model. At each position of the tested sequences, we replaced a nucleotide with a different one in all the sequences and computed their average score change. The rows represent nucleotides that are used for replacement and the columns show different positions inside the regulatory regions. If the new score on average increases, it is represented by a red-colored square. Decreasing the score is shown by using a blue-colored square. The intensity of color is proportional to the effect of substitution on the score. Since core promoters are very diverse, if we draw a mutation map for all the sequences, we cannot capture all the information from our model. To alleviate this, we clustered the mutation matrices based on their similarity (Frobenius norm) into several clusters using the k-means algorithm with k = 10.

Analysis of the mutation maps showed that promoters have higher GC content than enhancers and conserved *C* and *A* nucleotides at positions -1 and +1 respectively, which corresponds to the Initiator motif (**Table 2**). Enhancers do not have a conserved TSS nucleotide. However, clustering reveals that many of the enhancers have a *CCGACC* motif around the TSS. For promoters, the revealed motifs in the TSS region are *GGCATAC and GTCAGAC*.

Mutation/saliency maps and convolutional filter analysis are often used to understand deep learning models, but they cannot detect long-range interactions in the input sequence. We attempted to overcome this limitation by developing a novel technique to measure dependency between each pair of nucleotides - pair dependency map. **Figure 4a** shows the dependency between positions inside the core promoter region for the promoter sequences. Here every element of the matrix *V(i,j)* is the difference between two values: the change in the predicted score when removing the pair *(i, j)* compared to the sum of changes caused by removing *i* and *j* separately. Every input sequence generates a symmetric matrix, and we draw the averaged matrix. The red color is used for positive elements of *V*, and the blue color is for negative elements. Intensity shows the absolute value of the element.

**Figure 4.**
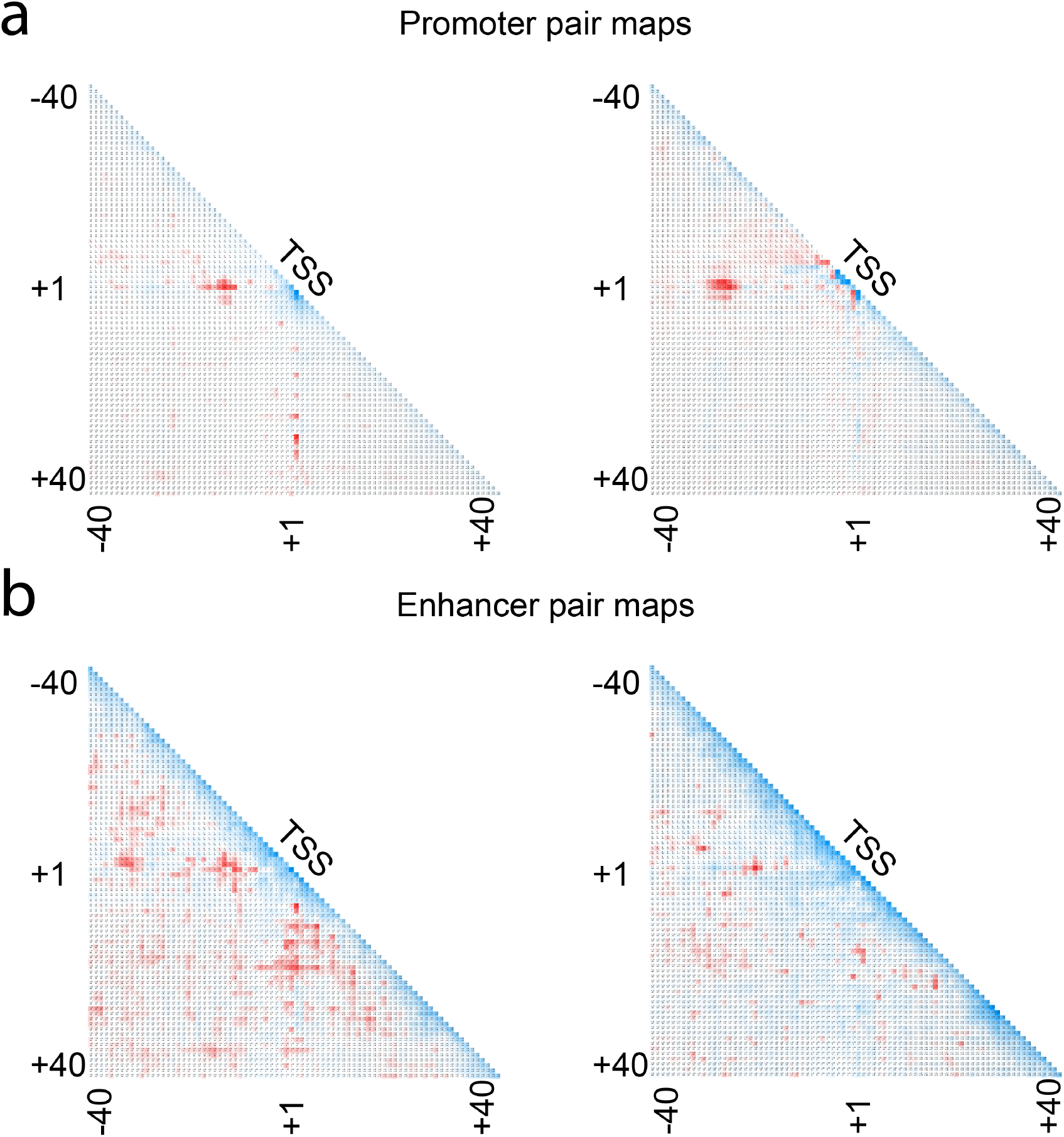
Pair dependency map reveals long-range interactions captured by our model. The red color means interaction, the blue color shows correlation, and the white color represents that the positions are independent. The results are clustered into ten groups using k-means algorithm and the two biggest groups are shown. **a.** Promoter pair maps show relationships between conserved promoter elements, which have much stronger interactions than other pairs. They include TSS interactions with TATA and BRE elements in the upstream region, DCE and DPE elements in the downstream region. **b.** Dependencies captured by our model for predicting enhancers are spread out in the [−40: +40] region. Unlike promoter sequences, there are no strong short-range interactions in the TSS region.

The pair maps for the enhancer sequences are shown in **Figure 4b**. As shown in previous experiments, interactions between TSS and the TATA-box are very important for the promoter sequences, while interactions in enhancer sequences are more uniformly distributed. This is especially easy to see after performing the k-means clustering of the matrices before averaging.

The proposed pair map analysis reveals related positions. Unlike the single position mutation map analysis, if a position changes a score significantly but independent of other positions, pairs involving it will be shown with a light color. The pair dependency map strongly suggests that our model can capture long range dependencies between different elements in the regulatory sequences, and that the trained model does not simply detect conserved core promoter elements but also complex interactions between them. Based on the pair map idea, we created a tool that can test whether two given positions (or two regions) are independent according to our model, which may be useful for creating hypotheses such as if two transcription factors regulate a subset of promoters together. Using the described approach, we analyzed interaction between JUND and BATF TFs, which was reported in the previous studies^36,37^. Searching for their motifs inside the regulatory regions using FIMO^38^ with default parameters, the interaction between JUND and BATF was compared to the interaction of JUND with random locations of the same motif length. As expected, a strong relationship between the two binding motifs was detected (p = 3.04e-11, t-test).

## Discussion

We have demonstrated that by using a dynamic training set, it is possible to tackle the problem of genome-wide regulatory elements prediction. This is a very difficult problem because many non-regulatory regions are very similar to the true ones in terms of their nucleotide sequences. Moreover, the current collection of promoters and enhancers might be incomplete, and many of the unannotated predictions generated by our method can be related to unknown or alternative promoters. Unlike in common machine learning problems, prediction at the genome-wide scale is extremely unbalanced, which makes it difficult to achieve high sensitivity. Despite all these challenges, the proposed model achieves a good performance on the human genome and can be directly applied to genomes of other animals as well without significant loss of performance.

In this study, we have shown that the trained model can identify motifs and positions important for the activity of regulatory elements, which allows for further exploration of the promoter architecture. Furthermore, we identified long-range interactions in enhancer and core promoter regions using a pair dependency map. The next step would be to validate these findings using reporter assays by constructing synthetic promoters and enhancers. This would help to better understand transcriptional regulation and also design new regulatory sequences with desired strength.

To further improve the results, it is necessary to consider the cell type when predicting promoters and enhancers. It has been shown that promoter prediction can also be improved using extra information as input, for example chromatin data^39^ or possibly applying nonparametric methods as described and tested on promoter regions of a model dicot plant Arabidopsis thaliana^40^. However, the approach described in this paper has the advantage that it is very general. It will be straightforward to apply it to many different organisms where additional information like chromatin profiles might not be available.

## METHODS

### Data sets used

The data we used for training our models consist of 210250 promoters and 65423 enhancers downloaded from FANTOM5 website. Sequences of length 1001 bp were extracted from the human genome (GRCh37) centered around the TSSs. All the regulatory regions from chromosome 1 were used for testing, 90% of the remaining peaks were used for training, and the rest for validation to choose deep learning parameters and perform early stopping during training. GENCODE version 34 was used when picking candidates for reporter assay validation to avoid picking regions near any known gene. GWAS and Clinvar sets were downloaded from their respective websites on 2020-07-06. The motif analysis used PWMs from the JASPAR 2020 version.

### Negative set generation

One of the reasons the reported performance in previous studies does not extrapolate to the whole human genome is because of an inadequately chosen set of negative sequences^15^. Often the negative set consists of random genomic sequences which are very different from the regulatory regions or sequences from specific positions (e.g., fixed distance away from TSS) which introduces bias. To tackle this problem, we used an iterative approach that updates the set of non-promoter sequences used in the training set based on the false positive errors made in the previous iteration^41^. By including difficult non-promoter sequences in the training set, the predictor is forced to learn promoter patterns to rule out such sequences. This scheme allowed us to achieve high sensitivity while keeping the number of false positive predictions low.

**Supplementary Figure 1** illustrates that without using these difficult negatives, the model mostly uses the GC content to make a prediction.

When a difficult negative set is obtained, the neural network struggles to discriminate provided positive and negative sequences. This can result in over-fitting, when the model simply memorizes some difficult negatives despite the regularization methods that are used. To avoid this issue, we have developed a new regularization technique. During each epoch, we added a small shift to the negative sequences, moving them upstream or downstream from the initial positions by a random distance, see **Supplementary Figure 2**. This virtually makes the negative set very large and impossible to memorize. The neural network can only learn very general patterns to rule out the negatives. We have found that this specific technique is superior to any other changes that one can do to the negative set. If parts of DNA sequence are manually changed, it is very easy for the neural network to detect such alterations, e.g., replacement, insertions, mirroring, generation of a new DNA sequence with specific nucleotide frequency. When such alterations are used, the network is trained to distinguish the original DNA sequence from an altered one, instead of discriminating promoters from non-promoters.

### Genome-wide prediction regulatory elements prediction

Since it is difficult to test every possible TSS location in the genome, we used a two-fold prediction procedure. One model is used to scan each chromosome using a sliding window approach. Because of the random shift added to both positive and negative sequences during the training, the input for the scan model does not need to be centered perfectly around the TSS. This allowed us to use a relatively large step for the sliding window, 50 bp. When the model outputs values above the threshold (0.5), the 100 bp region centered around the current position is scanned using position specific second model with the step equal to one. Predictions of the second model with scores higher than 0.5 are sorted and filtered based on the distance between them. The remaining results are output by the method. The models are trained to detect regulatory elements on both strands without explicitly performing reverse complement. The strand is decided by a special model, which decides the direction of transcription: either positive (+) or negative (-) for promoters and both (.) for enhancers. This design decision was made to make genome-wide TSS identification faster. During the evaluation, all the predictions more than 500 bp away from a known CAGE peak were considered as FPs. The margin for error is rather big to deal with the alternative promoters for the same gene, since the TSSs can be located a large distance from each other^42^.

### Deep learning architecture

The deep learning architecture is based on deep residual networks^43^. The network consists of 5 residual blocks followed by a Softmax layer, **Supplementary Figure 3**. To avoid the problem of vanishing gradient, we employed batch normalization^44^. The activation function used throughout the model is Leaky ReLU^45^. Weight decay and dropout are used to improve the generalization capability of the model. Weight decay effectively limits the number of free parameters in the model to avoid over-fitting. Introducing weight decay makes it possible to regularize the cost function by penalizing large weights. The main idea of dropout is to randomly set some nodes of the neural network to zero during training to prevent co-dependency among them. During the training, the dropout for the feature vector with keep probability of 0.5 is used. The Adam optimization algorithm is used to train the weights^46^, which is an improved version of stochastic gradient descent. TensorFlow^47^ is used as the framework to construct the deep neural network. The training was performed on a workstation with four NVIDIA Quadro RTX 6000 GPUs and took about 5 days to complete.

### Reporter assays

We used reporter assays to validate the candidate sequences. Double strand DNA cassette was constructed by annealing the DNA oligo that has SpeI site/I-CeuI site and the DNA oligo that has BamHII site/I-SceI site. Specific DNA primers were designed for amplification of the test candidates. Using these primers and Human Genomic DNA as template, PCR was performed with KOD-Plus-Neo. After QC with electrophoresis, PCR products (insert DNAs) were digested by I-CeuI/I-SceI. We selected and prepared human GAPDH promoter region as positive control and random sequences (backbone of pMCS-Cypridina Luc vector (201 bases long)) as negative control. The prepared vector and insert DNA were ligated using Rapid DNA ligation Kit, and One Shot TOP10 Chemically Competent E. coli was transformed by the ligated vector. These transformed E. coli cells were cultured in large scale and assay vectors were extracted using QIAGEN Plasmid Mini Kit. For quality checking, PCR was performed with assay vector and primers for checking. After this, electrophoresis was performed to confirm the presence of the desired insert DNA fragment in the assay vector.

Until the day before doing assay, HEK293T cells were pre-cultured. At the day before doing assay, pre-cultured HEK293T cells were spread on 96-well plate and incubated at 37°C CO2 5% for 16-24h. After incubation, assay vector was transfected into HEK293T cells using Turbofect Transfection Reagent according to the kit protocol. After incubation, Cells were lysed using Cell lysis buffer included in Pierce Cypridina-Firefly Luciferase Dual Assay Kit. Prepared reagent mixture containing D-Luciferin was added to the cell lysate and measurement of luminescence from luciferin-luciferase reaction was performed by luminometer. Detailed description of the experimental setup is provided in the **Supplementary Appendix A**.

### Alternative methods

We have obtained Basenji from the GitHub repository and followed the provided guide for peak prediction (https://github.com/calico/basenji/tree/master/manuscripts/basset). The data were replaced with two of our bed files for promoters and enhancers. After generating the data, we modified the target neurons number, setting it to 2. The training took 18 epochs and was stopped automatically after the validation AUROC did not improve (0.75301). The trained model was then applied to chromosome 1 with the default test stride of 192 using provided script: basenji_predict_bed.py. EP3 and PromPredict do not require extra training, which is why we applied them directly on both chromosome 1 and its reverse complement.

### Synthetic promoters analysis

The synthetic sequences were inserted into the same genomic background as in the original publication before using them as an input for our model. To undo the effect of Softmax which makes all the output values very close to 0 or 1, our predicted score was adjusted as follows: log (1+*Score*/(1 – *Score*)).

## Data and code availability

Trained models, code to generate them, and code for analysis and figures described in this study are available at the following GitHub repository: https://github.com/umarov90/ReFeaFi. All the data used in training and validation are publicly available through databases referenced above.

## Acknowledgements

We would like to thank Andrew Tae-Jun Kwon and Bogumil Kaczkowski for insightful comments on the manuscript.

## Author Contributions

EA conceived the study and supervised the research together with XG. RU performed the research and drafted the manuscript together with EA. YL helped to validate the method. TA and ST performed the reporter assay.

