## Supplementary information for "ReFeaFi: Genome-wide prediction of regulatory elements driving transcription initiation"

| Species | Genome size | Number of robust CAGE peaks | Peaks per 1 Mb |
|---|---|---|---|
| Human | 3095693983 | 201802 | 65 |
| Mouse | 2654911517 | 158966 | 60 |
| Chicken | 1018836108 | 31863 | 31 |
| Rat | 2782012602 | 28325 | 10 |
| Dog | 2327650711 | 23147 | 10 |
| Monkey | 2824226272 | 25869 | 9 |

**Table 1.** Number of CAGE peaks for different species.

| ID | Chr | Position | Closest peak | Strand | Conservation | TATA | Inr | GC% |
|---|---|---|---|---|---|---|---|---|
| TS1 | chr20 | 55500504 | 62721 | + | 0.95 | 0 | 0 | 59 |
| TS2 | chr17 | 37719879 | 43991 | - | 0.99 | 0 | 0 | 59 |
| TS3 | chr16 | 51147623 | 35748 | + | 0.94 | 0 | 0 | 52 |
| TS4 | chr14 | 105310505 | 12977 | - | 0.00 | 0 | 0 | 55 |
| TS5 | chr2 | 121344650 | 23142 | + | 0.00 | 0 | 0 | 72 |
| TS6 | chr6 | 99797621 | 75121 | + | 0.40 | 1 | 0 | 72 |
| TS7 | chr3 | 53528516 | 146947 | - | 0.22 | 0 | 1 | 82 |
| TS8 | chr19 | 44913646 | 52505 | + | 0.00 | 1 | 1 | 62 |
| TS9 | chr7 | 1315350 | 14836 | + | 1.00 | 0 | 0 | 52 |

| | | | | | | | | |
|---|---|---|---|---|---|---|---|---|
| TS10 | chr4 | 17135852 | 76196 | + | 0.93 | 0 | 0 | 41 |
| TS11 | chr6 | 45709523 | 13093 | + | 0.89 | 0 | 0 | 45 |
| TS12 | chr16 | 88186706 | 41694 | + | 0.00 | 0 | 0 | 63 |
| TS13 | chr1 | 229039512 | 35035 | + | 0.00 | 0 | 0 | 49 |
| TS14 | chr4 | 189167477 | 23262 | + | 0.00 | 0 | 0 | 57 |
| TS15 | chr13 | 25572503 | 49119 | + | 0.03 | 1 | 0 | 52 |
| TS16 | chr14 | 97165660 | 19170 | + | 0.01 | 0 | 1 | 50 |
| TS17 | chr9 | 122573287 | 27611 | + | 0.47 | 1 | 1 | 37 |
| TS18 | chr4 | 58581891 | 604807 | - | 0.08 | 0 | 0 | 40 |
| TS19 | chr2 | 105224916 | 114521 | - | 0.06 | 0 | 1 | 45 |
| TS20 | chr2 | 21882925 | 590849 | - | 0.36 | 0 | 1 | 43 |

**Table 2.** Predicted regulatory sequences that were chosen for reporter assay validation.

| Promoters | Enhancers |
|---|---|
| 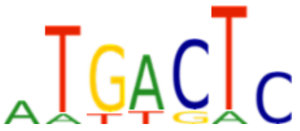 | 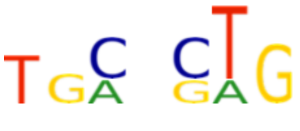 |
| 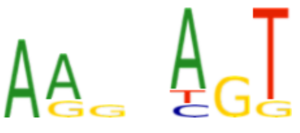 | 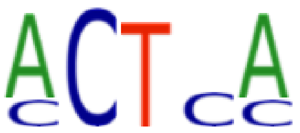 |
| 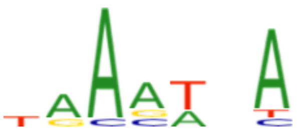 | 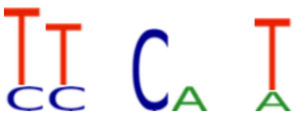 |
| 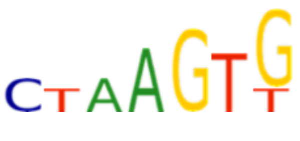 | 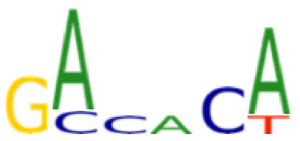 |
| 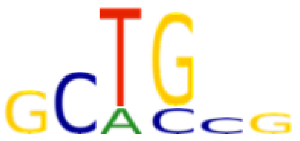 | 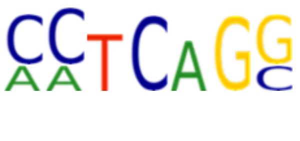 |
| 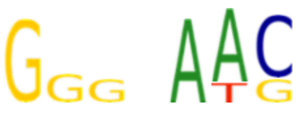 | 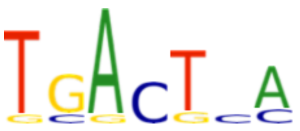 |

**Table 3.** Most important motifs identified by our model outside the core promoter region which do not match JASPAR motifs.

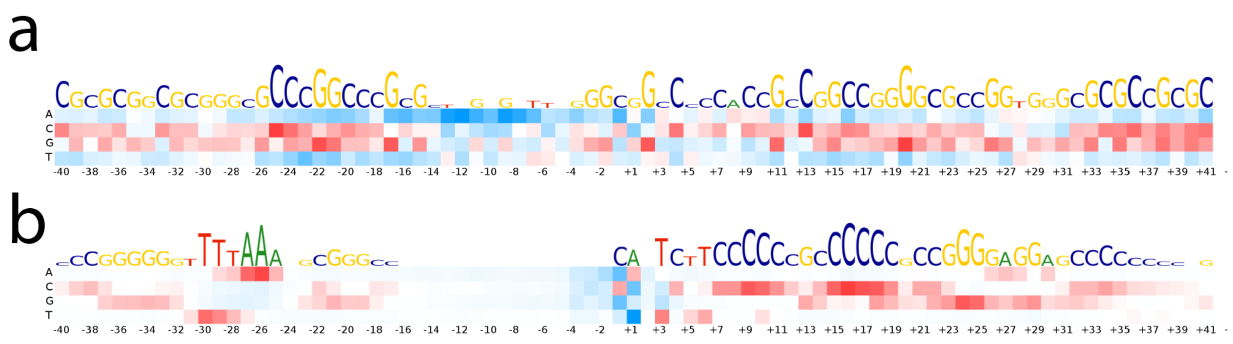

**Figure 1.** Difference between random negatives model and hard negatives model in terms of their mutation map. **a.** Random negatives model. **b.** Difficult negatives model.

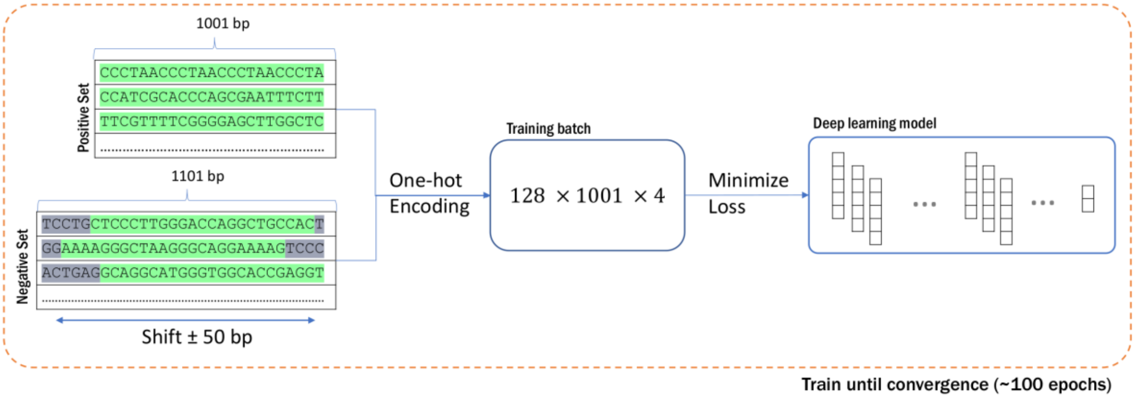


**Figure 2.** Procedure of shifting negative sequences, which virtually increases the negative set size and prevents overfitting.

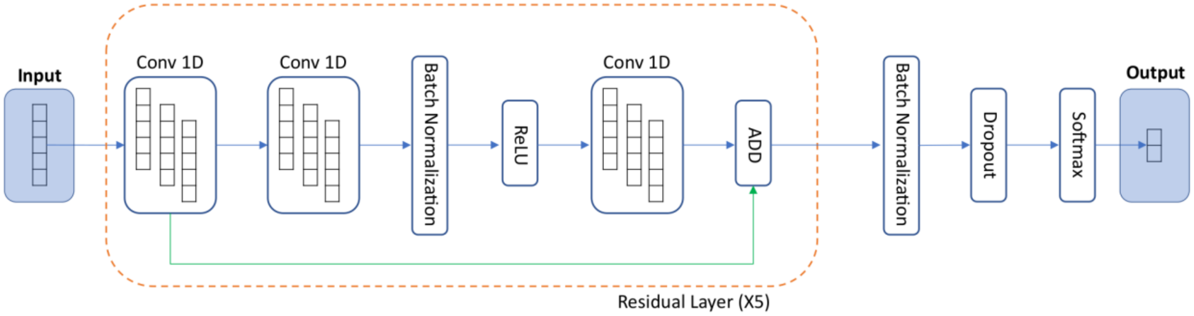


**Figure 3.** Deep learning architecture that was used in building our model. The model employs CNN with residual connections which is followed by a softmax layer.

### Appendix A

##### Vector modification, promoter candidate fragment preparation and assay vector preparation

pMCS-Cypridina Luc vector (Cat#16149, Thermo Fisher SCIENTIFIC) has eight cloning sites and all of them can be cut by six base recognition restriction enzyme. But in that case, some desired promoter candidate region may have that recognition site. It is cut in the middle of DNA fragment. It is inconvenient for ligating. To perform ligation efficiently, we decided to modify the vector to have a Homing endonuclease site that recognizes rare sequences.

Double strand DNA cassette was constructed by annealing the DNA oligo that has SpeI site/I-CeuI site and the DNA oligo that has BamHII site/I-SceI site (DNA oligos were synthesized by Eurofin Genomics). And next, its DNA cassette was digested with SpeI (Cat#R0133S, New England Biolabs)/BamHI-HF (Cat#R3136S, New England Biolabs). On the other hand, pMCS-Cypridina Luc vector (Cat#16149, Thermo Fisher SCIENTIFIC) was digested with SpeI/BamHI-HF and both 5'-ends of vector were dephosphorylated with FastAP Thermosensitive Alkaline Phosphatase (Cat#EF0654, Thermo Fisher SCIENTIFIC).

DNA cassette after digestion with SpeI/BamHI and the vector after digestion with SpeI/BamHI were ligated using Rapid DNA ligation Kit (Cat# K1422, Thermo Fisher SCIENTIFIC). Next, One Shot TOP10 Chemically Competent E. coli (Cat# C404006, Thermo Fisher SCIENTIFIC) was transformed by the ligated vector. These transformed E. coli cells were cultured in large scale and modified vector was extracted using QIAGEN Plasmid Mini Kit (Cat# 12123, QIAGEN). It is named “pMCS-Cypridina Luc wCST”. pMSC-Cypridina Luc wCST was digested with homing endonuclease I-CeuI

(Cat#R0699S, New England Biolabs) and I-SceI (Cat#R0694S, New England Biolabs), and both 5'-ends of vector were dephosphorylated with FastAP Thermosensitive Alkaline Phosphatase.

Specific DNA primers were designed for amplification of promoter candidate (DNA primers were synthesized by Eurofin Genomics). Using these primers and Human Genomic DNA (Cat# G1471, Promega) as template, PCR was performed with KOD-Plus-Neo (Cat#KOD-40, TOYOBO). Reaction conditions were followings.  Basically 2 step PCR (94℃ 2min->[98℃ 10sec->68℃ 30sec] x 35cycles->68℃ 10min) was performed. PCR of the sample that appeared multi-bands were performed with step-down PCR protocol (94℃ 2min->[98℃ 10sec->68℃ 30sec] x 35cycles->68℃ 10min). As for PCR for the samples in the result of no amplification, 3step PCR protocol (94℃ 2min->[98℃ 10sec->74℃ 30sec] x 5cycles->[98℃ 10sec->72℃ 30sec] x 5cycles->[98℃ 10sec->70℃ 30sec] x 5cycles->[98℃ 10sec->68℃ 30sec] x 25cycles-> 68℃ 10min) was used. After QC with electrophoresis, PCR products (insert DNAs) were digested by I-CeuI/I-SceI. We selected and prepared human GAPDH promoter region as positive control and random sequence (backbone of pMCS-Cypridina Luc vector (201 bases length)) as negative control.

The above prepared vector and insert DNA were ligated using Rapid DNA ligation Kit, and One Shot TOP10 Chemically Competent E. coli was transformed by the ligated vector. These transformed E. coli cells were cultured in large scale and assay vectors were extracted using QIAGEN Plasmid Mini Kit. For quality checking, PCR was performed with assay vector and primers for checking. After this, electrophoresis was performed to confirm the presence of the desired insert DNA fragment in the assay vector. In addition to the self-made control vector, pCMV-Cypridina Luc (Cat # 16150, Thermo Fisher SCIENTIFIC) purchased as a positive control and pMCS-Cypridina Luc w CST with no insert DNA were prepared as a negative control.

**Assay vector transfection and luciferase assay**

Until the day before doing assay, HEK293T cells were pre-cultured (DMEM (Cat#11965-092, Gibco) + 10% FBS (Cat# 26140-079, Gibco)). At the day before doing assay, pre-cultured HEK293T cells were spread on 96-well plate (Cat#167008, Thermo Fisher SCIENTIFIC), at the number of $1x10^4$ /well in DMEM+10%FBS medium and was incubated at 37℃ $CO_2$ 5% for 16-24h. After incubation, assay vector was transfected into HEK293T cells using Turbofect Transfection Reagent (Cat#R0533, Thermo Fisher SCIENTIFIC) according to the kit protocol. To correct the difference of luminescence intensity caused by the difference of transfection efficiency and cell number, pCMV-Red Firefly Luc vector (Cat#16156, Thermo Fisher SCIENTIFIC) was co-transfected together. Vector-transfected HEK293T cells were incubated 37℃ $CO_2$ 5% for 24h. After incubation, Cells were lysed using Cell lysis buffer included in Pierce Cypridina-Firefly Luciferase Dual Assay Kit (Cat# 16183, Thermo Fisher SCIENTIFIC). According to the kit protocol, prepared reagent mixture containing D-Luciferin were added to the cell lysate and measurement of luminescence from luciferin-luciferase reaction was performed by luminometer (2030 ARVO X, Perkin Elmer).

**Original vector for assay (before modification)**

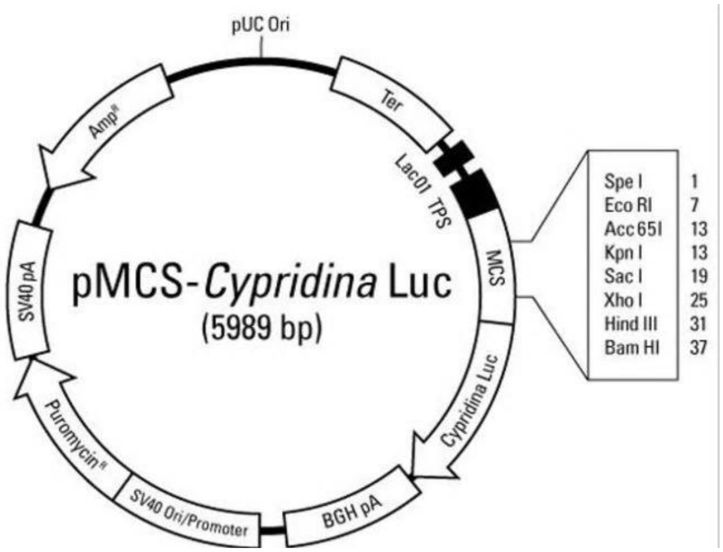


**Assay v[illegible]**

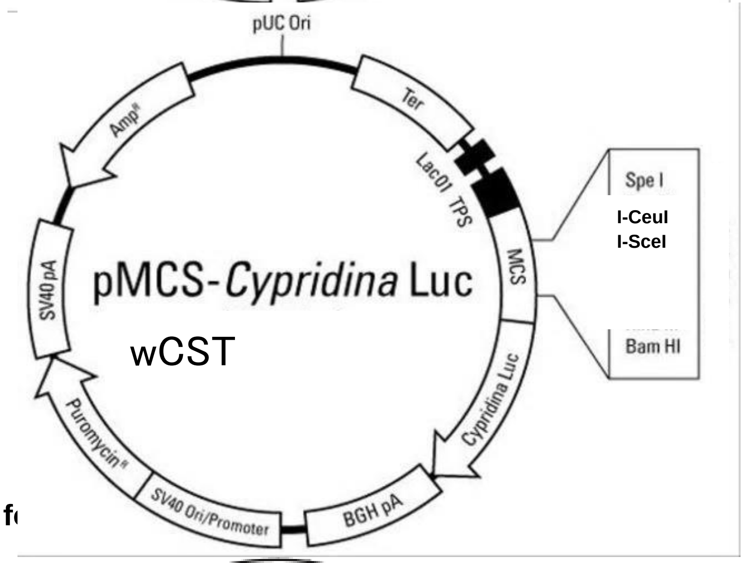


**Vector f[illegible]**

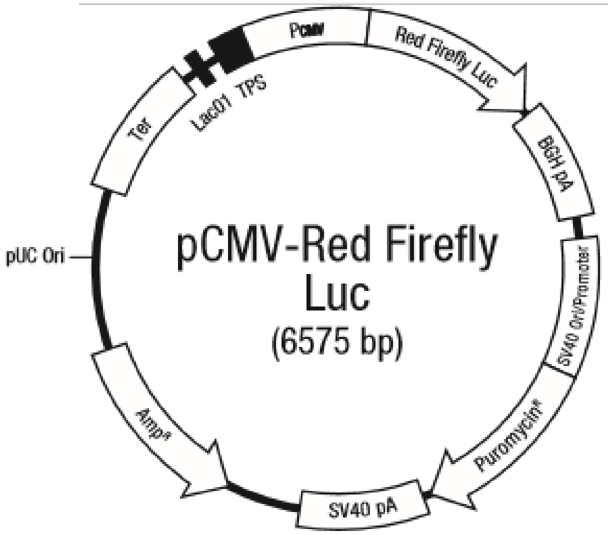

### Outline of experiment (vector modification - construction of assay vector)

- cassette
  SpeI = I-CeuI ======== I-SceI = BamHI
- vector
- insert

DNA cassette after annealing

```
5'-GGGGAˇCTAGTTAACTATAACGGTCCTAAˇGGTAGCGAAGAGAGAGAGAGAGAGAGAGATAGGGATAAˇCAGGGTAATGˇGATCCCCCC-3'
3'-CCCCTGATC-AATTGATATTGCCAG-GATTCCATCGCTTCTCTCTCTCTCTCTCTCTCTATCCC-TATTGTCCCATTACCTAG-GGGGG-5'
```

SpeI/BamHI treatment for DNA cassette

```
5'-------CTAGTTAACTATAACGGTCCTAAˇGGTAGCGAAGAGAGAGAGAGAGAGAGAGATAGGGATAAˇCAGGGTAATG---------3'
3'-----------AATTGATATTGCCAG-GATTCCATCGCTTCTCTCTCTCTCTCTCTCTCTATCCC-TATTGTCCCATTACCTAG-----5'
```

SpeI/BamHI treatment &dephosphorylation by Alkaline phosphatase

```
5'-NNNNA--------------------------------------------------------------------------GATCCNNNN-3'
3'-NNNNTGATC--------------------------------------------------------------------------GNNNN-5'
```

Ligation with DNA cassette with original vector

```
5'-NNNNACTAGTTAACTATAACGGTCCTAAˇGGTAGCGAAGAGAGAGAGAGAGAGAGAGATAGGGATAAˇCAGGGTAATGGATCCNNNN-3'
3'-NNNNTGATCAATTGATATTGCCAG-GATTCCATCGCTTCTCTCTCTCTCTCTCTCTCTATCCC-TATTGTCCCATTACCTAGGNNNN-5'
```

I-CeuI/I-SceI treatment & dephosphorylation by Alkaline phosphatase for modified vector

```
5'-NNNNACTAGTTAACTATAACGGTCCTAA----------------------------------------CAGGGTAATGGATCCNNNN-3'
3'-NNNNTGATCAATTGATATTGCCAG------------------------------------------TATTGTCCCATTACCTAGGNNNN-5'
```

DNA fragment synthesis of desired promoter candidate region by PCR

```
5'-GGGGTAACTATAACGGTCCTAAˇGGTAGCGAANNNNNNNNNNTAGGGATAAˇCAGGGTAATCCCC-3'
3'-CCCCATTGATATTGCCAG-GATTCCATCGCTTNNNNNNNNNNATCCC-TATTGTCCCATTAGGGG-5'
```

I-CeuI/I-SceI treatment for insert DNA fragment

```
5'------------------------GGTAGCGAANNNNNNNNNNTAGGGATAA--------------3'
3'-------------------GATTCCATCGCTTNNNNNNNNNNATCCC-----------------5'
```

Ligation with insert DNA fragment with modified vector

```
5'-NNNNACTAGTTAACTATAACGGTCCTAAGGTAGCGAANNNNNNNNNNTAGGGATAACAGGGTAATGGATCCNNNN-3'
3'-NNNNTGATCAATTGATATTGCCAGGATTCCATCGCTTNNNNNNNNNNATCCCTATTGTCCCATTACCTAGGNNNN-5'
```

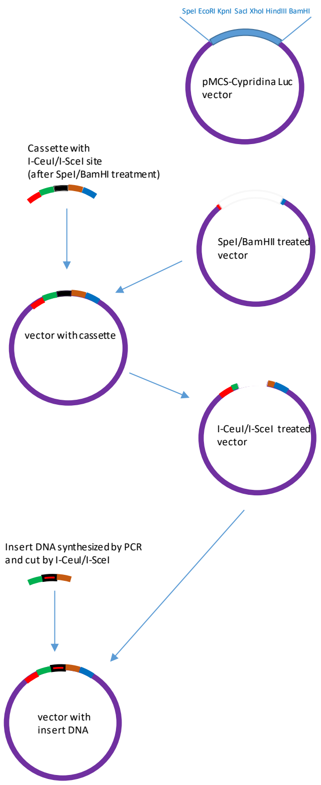

**Assay vector transfection to HEK293T**

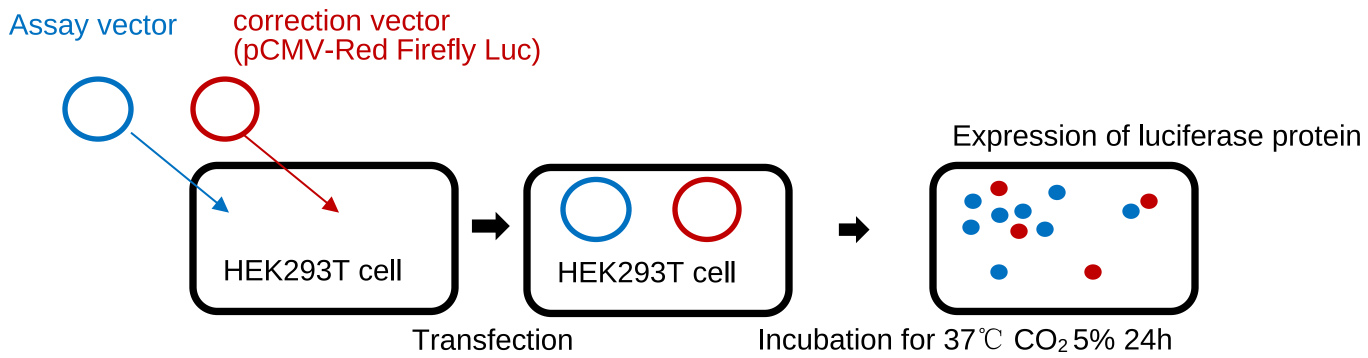


**Luciferase assay**

(Referred from Thermo Fisher SCIENTIFIC)

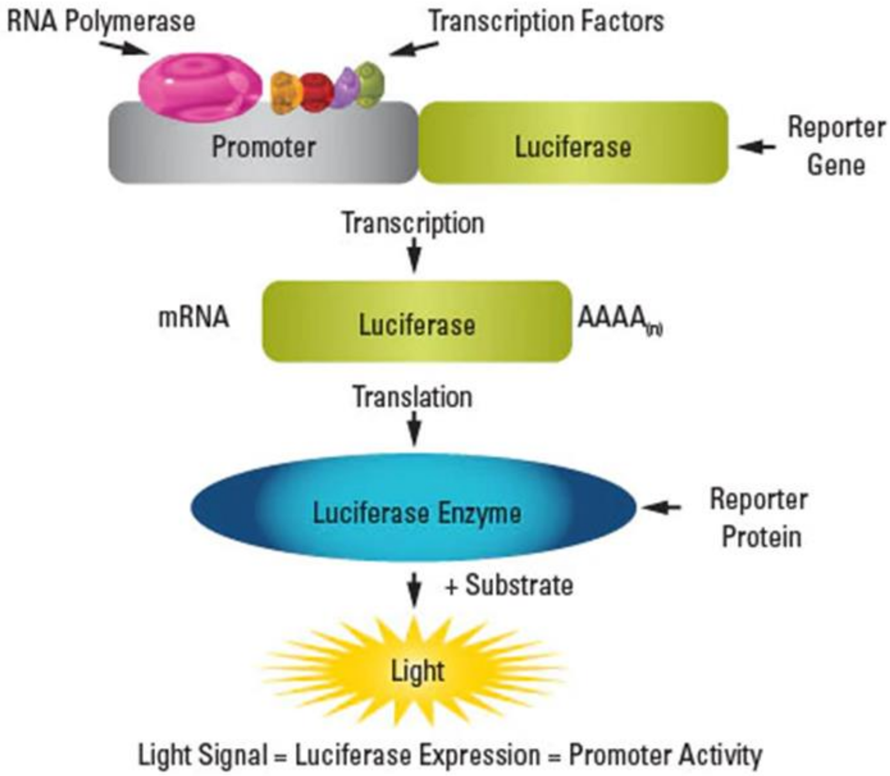
